# Method for Spatially Confined Stimulation by Retinal Scanning Displays: Are Blind Spots Really Blind?

**DOI:** 10.1101/2023.10.03.560249

**Authors:** Amir-Vala Tavakoli, Takanobu Omata, Teppei Imamura, Ryo Ogawa, Masanori Iwasaki, Jesus del Rio Salgado, Iyla Rossi, Shao-Min Hung, Daw-An Wu, Shinsuke Shimojo

## Abstract

Conventional displays, such as CRTs and LCDs, are commonly used to present and manipulate visual stimuli in many vision science studies. However, such displays pose difficulties when the goal is to constrain stimulation to specific retinal locations, so as to isolate their contribution to downstream effects. Scattered and refracted light can stimulate unintended areas, contaminating the outcomes. In this investigation, we evaluate a laser-based retinal scanning display for its ability to map and exclusively stimulate the retinal blind spot. This application stimulates melanopsin receptors inside the blind spot, separately from rod and cone photoreceptors, which lie outside the blind spot. This paper covers three experiments: 1. The initial exploratory observations; 2. the examination of pupillary responses, and; 3. the inference of implicit visual perception arising from the blind spot. We aimed to simultaneously validate our technical methods, while increasing our understanding of the role of melanopsin-expressing intrinsically photosensitive retinal ganglion cells (ipRGCs) in vision and implicit visual perception.

## INTRODUCTION

Retinal scanning displays can draw images onto the retina directly through the pupil by scanning laser beams across the surface. The displays are used for various industrial applications such as augmented reality and as solutions for low vision patients, but we would like to propose that the displays could be advantageous for a variety of vision studies. Here, we report our work in adapting the display to a study combining psychophysical and neural recordings using electroencephalography (EEG).

Specifically, we will address the following issues: ipRGCs are a third type of photosensitive cell in the human retina whose function is less understood than that of the classical rod and cone photoreceptors. Unlike rods and cones, ipRGCs express their photopigment even in the blind spot. One way of studying this visual pathway in isolation, then, is to confine experimental stimuli to the blind spot. Could a retinal scanning display improve data quality and potentially modify current knowledge of the ipRGCs, by leveraging the specific advantages of the display’s laser and avoiding the refraction of light inside the eye? Could this increased anatomical specificity be used to investigate the psychological and neural functions of those ipRGCs within the blind spot? What are the potential advantages and disadvantages from a technical perspective? Using a laser scanning display to replicate previous reports would be worthwhile to evaluate the possibility that their results may have been contaminated by artifacts from refracted light. Differences in outcomes between conventional and laser stimulation would indicate the importance of using laser stimulation in this type of study.

## 1. Brief review: Human studies of ipRGC

For most of the last 150 years, the scientific understanding held that mammalian vision is mediated by phototransduction in two types of photoreceptors, rods and cones (Graham & Wong, 2016; Sanes & Masland, 2015). It was not until 2002 that a new “photoreceptive net” was identified in mice (Provencio et al., 2002), based on a new kind of light-reactive protein called melanopsin (Provencio et al., 1998). Because the gene for melanopsin photopigment is found throughout the mammalian class, it was not long before its presence was confirmed in other mammals, including humans (Bito & Turansky, 1975; Lau et al., 1992; Tu et al., 2004).

The traditional rod and cone receptors pass their signals inward across layers of retinal cells, ultimately to the innermost layer of retinal ganglion cells (RGCs; Mure, 2021; Schmidt & Kofuji, 2009). Within the dendritic network of RGCs, some RGCs are dotted with melanopsin proteins that function as light detectors (Hattar et al., 2002; Panda et al., 2002). The subset of RGCs expressing melanopsin were dubbed intrinsically photosensitive retinal ganglion cells, or ipRGCs (Berson et al., 2002). Their sensitivity in the light spectrum was investigated and reliably found in the range of 460-500 nm (Cajochen et al., 2005; Hattar et al., 2003; Lockley et al., 2006; Lucas et al., 2001; Takahashi et al., 1984). Melanopsin’s blue light sensitivity mediates the transduction of light conditions in our environment, modulating circadian functions, the central nervous system, and autonomic functions (Aranda & Schmidt, 2021; G. C. Brainard et al., 2001; Chellappa et al., 2016; Hattar et al., 2002, 2006; Klerman et al., 2022; LeGates et al., 2014; Lerner et al., 1959; Wiechmann et al., 1986).

Melanopsin is distributed widely in the retina, including in the optic disk, also known as the blind spot. This area is occupied entirely by the bundle of RGC axons passing out of the eye. As a consequence, there are no rod and cone photoreceptors in this area, only the melanopsin receptors expressed in the ipRGC axons. Precise stimulation of this area would elicit melanopsin activity, theoretically in the absence of all other inputs, providing an opportunity for scientists to selectively explore the function of the melanopsin/ipRGC pathway. ipRGCs respond not only to their own melanopsin receptors, but also in response to signals relayed to them from rod and cone receptors (Mure, 2021). Thus, investigation of interactions between photoreceptor types is also possible. Controlled stimulation of both populations can reveal how the output of ipRGCs modulates the activity of surrounding cones and rods.

## 2. Approaches of previous studies

In animal and in vitro studies, it is possible to perform genetic and biochemical manipulations to either eliminate or deactivate rods and cones. This leaves only the melanopsin on the ipRGCs to respond to light, so that they can be studied in isolation. These manipulations are not available in human studies, mainly due to the invasive nature of those techniques. While investigation of the human retina prompted development of a variety of alternative approaches, varying in their respective contributions, it has historically been necessary to scatter light across the retina in the process of presenting visual stimuli (Kim et al., 2021).

Studies have also been conducted on blind patients, with no conscious perception of light, wherein light has been reported to suppress both melatonin secretion (Czeisler et al., 1995; Hull et al., 2018) and remaining pupillary light reflexes (Charng et al., 2017). Zaidi et al. showed the existence of a similar system of inner retinal photoreception in rodless and coneless humans, similar to that found in some mammals, through experiments using patients with rod and cone dystrophies (Zaidi et al. 2007). However, these studies focus on blind patients so the results may differ from the response of typically developing humans.

As an alternative, a few papers have taken the approach of stimulating the blind spot (Miyamoto & Murakami, 2015; Saito et al., 2018). This is a particularly elegant approach, because this part of the retina is occupied by the optic nerve bundle exiting the eye, where there are no rod and cone receptors. This bundle consists of all the RGC axons projecting out of the eye and toward the brain. Because the melanopsin-containing ipRGCs are a subset of this bundle (Hattar et al., 2002), they can be stimulated by shining light into the blind spot. This is the one location where melanopsin inputs can be segregated, physically, from rod and cone inputs. The studies stimulating the blind spot are promising, but have thus far been done with traditional stimulation, e.g., light coming from computer monitors. This leaves the possibility that some of the light which is meant to stimulate the blind spot will scatter as it passes through the intraocular medium, traveling to other parts of the retina, and activating rods and cones. Thus, the results cannot be guaranteed to be based exclusively on ipRGC responses. In addition, these studies focused on colors and timescales not optimized for melanopsin properties (Gamlin et al., 2007; Miyamoto & Murakami, 2015; Saito et al., 2018). Melanopsin is sensitive to blue light, and responds quite slowly (Ecker et al., 2010; Schmidt & Kofuji, 2009; Wong, 2012; Zhao et al., 2014).

## 3. Our Approach

Here, we use a retinal scanning display. The displays consist of a small laser projector that draws images onto the retina, directly through the pupil, by laser beam scan using Micro Electro Mechanical Systems (MEMS; For a review see Lin et al., 2016). Retinal scanning displays are used for presenting visual images in augmented reality environments (e.g., Peillard et al., 2020), but they avail a set of important benefits to investigators of vision, for example resolving vergence-accommodation conflict. Laser projection in this way affords investigators a means of adopting the Maxwellian view, to project pure beams of light (Maxwell, 1860; Westheimer, 1966). The displays allow for rare precision in the presentation of visual stimuli, while still being noninvasive, an essential component of investigating non-clinical, human populations. The laser allows investigators to avoid optical artifacts induced by digital displays that broadly or non-specifically illuminate the retina of typically developing humans. By projecting the laser’s spatially precise beam over the inner retinal surface, onto the irregular boundaries of the blind spot, as well as the visual field, it is possible to investigate retinal functions specifically with respect to the corresponding gradient of diverse cell-types (Aranda & Schmidt, 2021). That is, it is possible to illuminate the retina with pure wavelengths of light, or to apply this same precision to ipRGCs within the blind spot. The unique properties of retinal scanning displays thus allow for a precise match between the stimulus and the ipRGCs in terms of spatial arrangement and frequency specificity.

The advantage of this method is to present collimated light through the pupil, minimizing the cross section subject to scatter, and focusing the presentation regardless of lens accommodation. We have chosen this topic of investigation (ipRGCs’ function in the blind spot) to highlight and to distinguish the advantages of the laser stimulator.

## 4. Purpose of this study

The purpose of this study was to develop a method for investigating the selective stimulation of melanopsin within the human retina. With the help of a retinal scanning display (RSD), we established a method of direct stimulation to the retina, and combined the retinal scanning display with psychophysics and EEG. In order to establish the method, we developed an original hardware set-up and blind spot mapping method. Then, we tested the method to find its problems through some initial exploratory experiments (Experiment 1: The initial exploratory observations) in order to check the feasibility. After that, we applied the designs of previous studies (Gamlin et al., 2007; Miyamoto & Murakami, 2015) to test the validity of our approach and find its advantages and disadvantages (Experiment 2: The examination of pupillary responses). We also sought to examine subliminal effects on visual attention, as evidenced by eye movements (Experiment 3: The inference of implicit visual perception arising from the blind spot). We aimed to simultaneously validate our method, while increasing our understanding of the role of melanopsin-expressing intrinsically photosensitive retinal ganglion cells (ipRGCs) in vision and implicit visual perception.

## 5. Methods Hardware set-up

We used a prototype retinal scanning display, which maintains the same principle as developed by Sony Group Cooperation (Akutsu et al., 2019). This prototype presents three different wavelengths of light, 445 nm, 522 nm, and 643 nm (blue, green, and red), using a MEMS and a holographic optical element (HOE) to scan the beams across the retina. Before beginning each experimental session, the laser was calibrated with a power meter to check for abnormalities in the output of the display and to confirm the laser’s output with regard to ipRGCs’ peak sensitivity, the instrument’s stability, and safety of intensities.

As shown in figures 1 and 2, the display and its baseplate were set on top of an optical bench. Additional optical mounts were added to the baseplate and the optical bench, to hold additional equipment and to provide mechanical stability: An eye tracking camera and infrared light source, which were partially disassembled from an Eyelink2 eye tracking system (SR Research, Canada); A fiducial fixation point, printed on cardboard-backed paper; A simple red laser pointer, attenuated with layers of neutral density filter, directed at the fixation target; A physical array of eye tracking calibration target stickers, affixed to a sheet held by a posterboard stand; A bracing structure that stabilized the chinrest. The chinrest (Takei Scientific Instruments, Japan) was customized by widening the width to allow other devices to be set in the limited space. Figures 2-4 show a participant during visual stimulation and the experimental setup.

**Figure 1.**
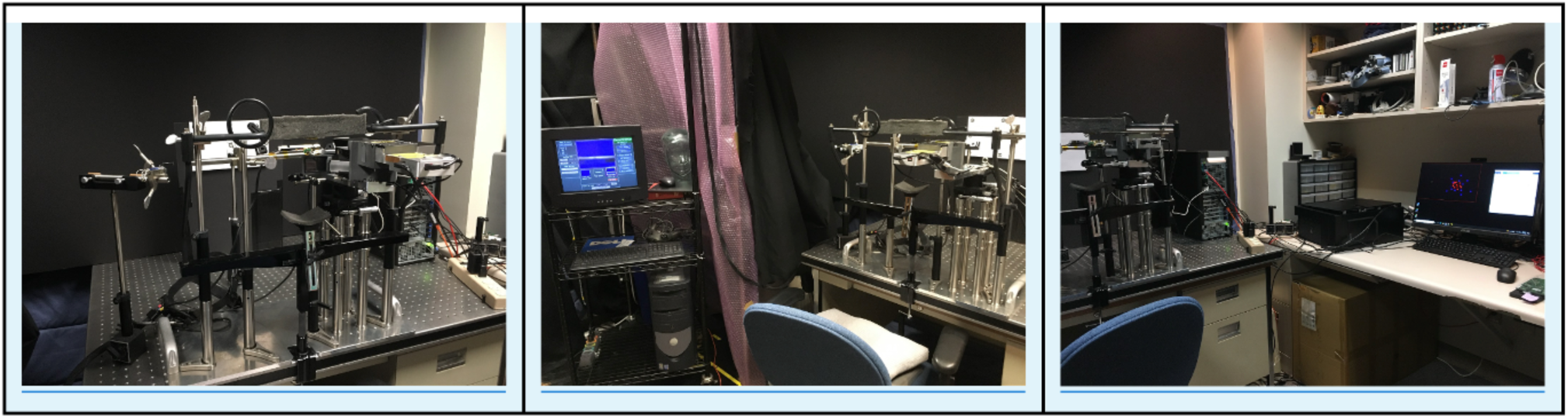
Experimental Room at California Institute of Technology

**Figure 2.**
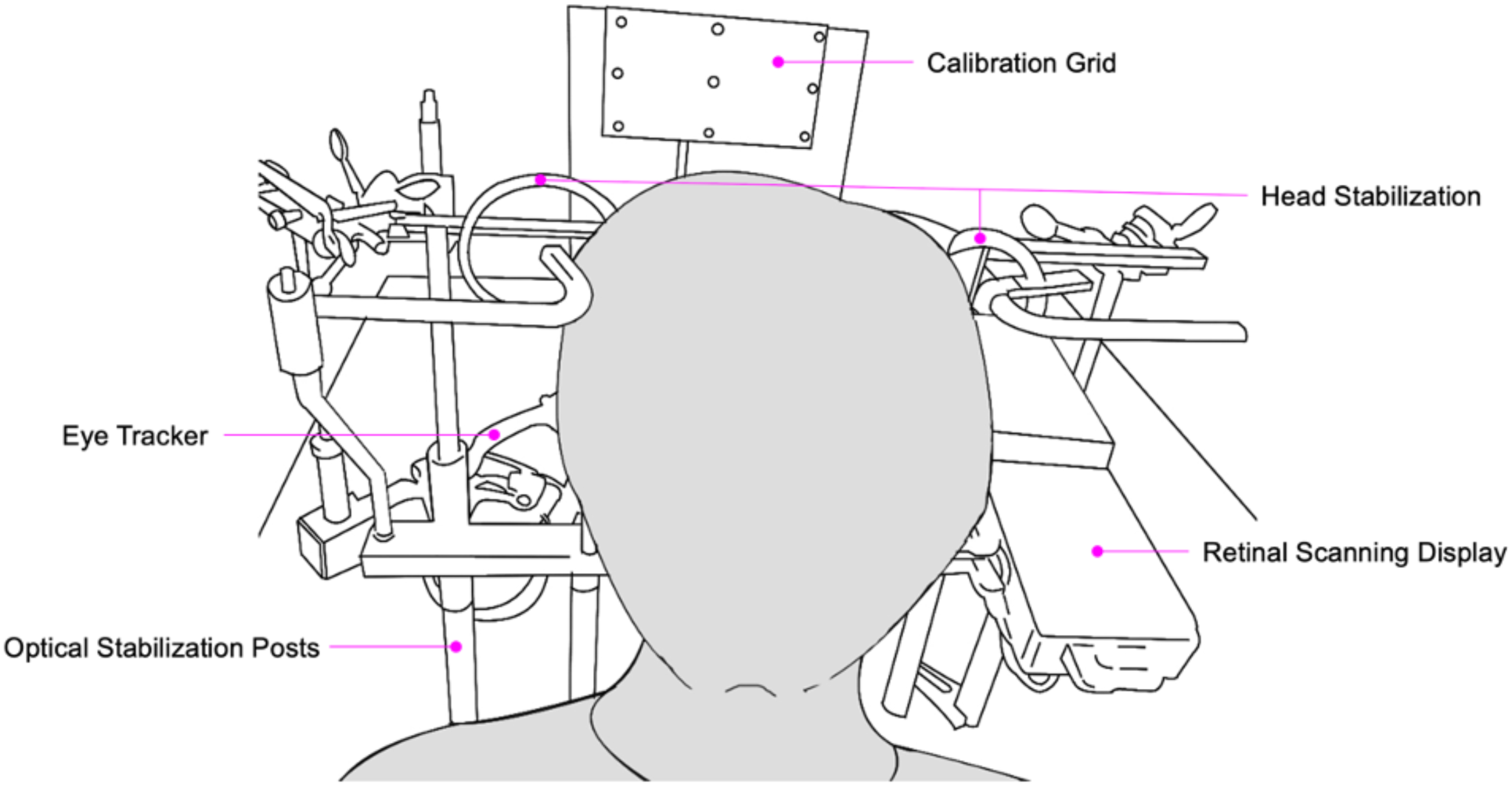
Schematic of participants’ interface with the experimental setup

**Figure 3.**
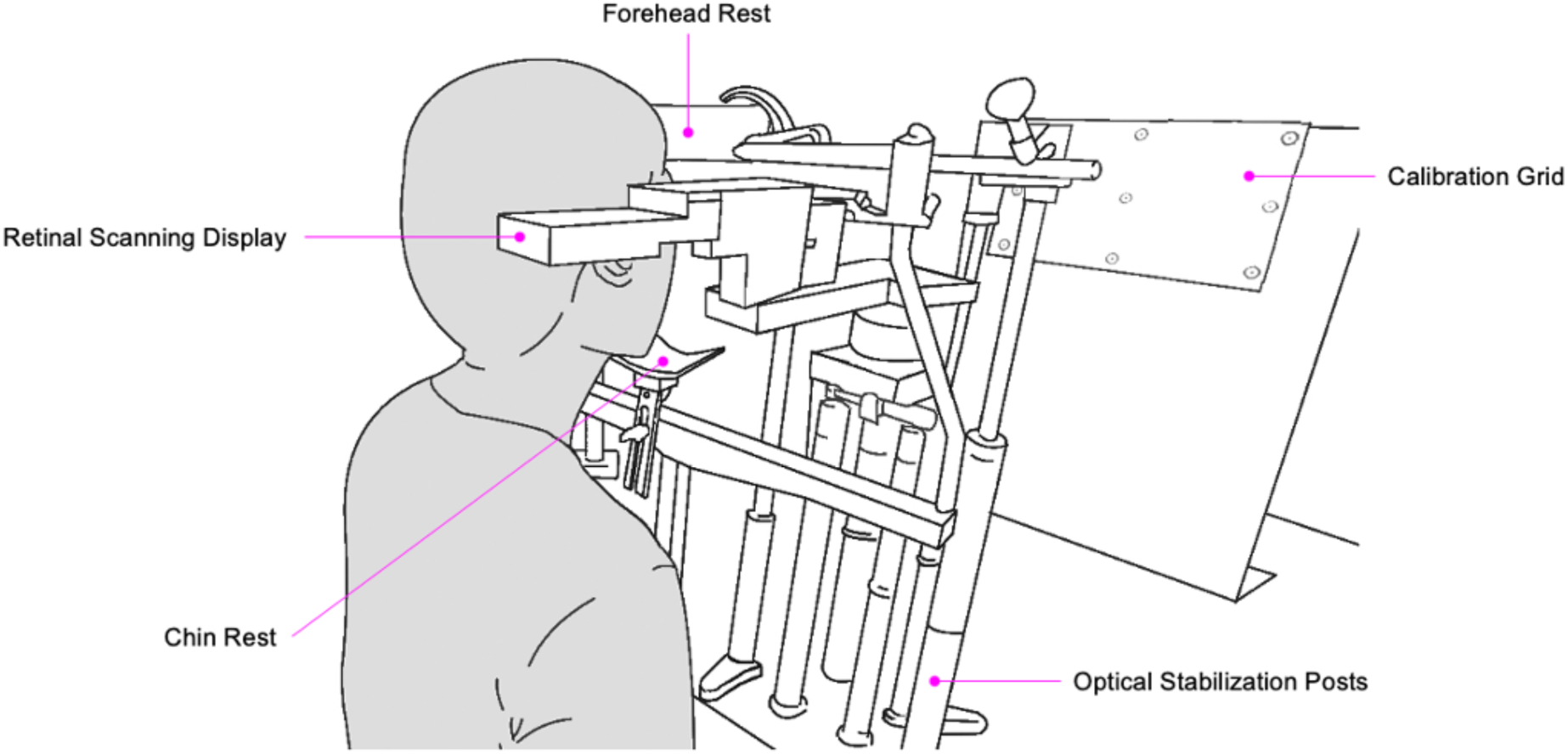
Schematic of experimental setup and visual stimulation

**Figure 4.**
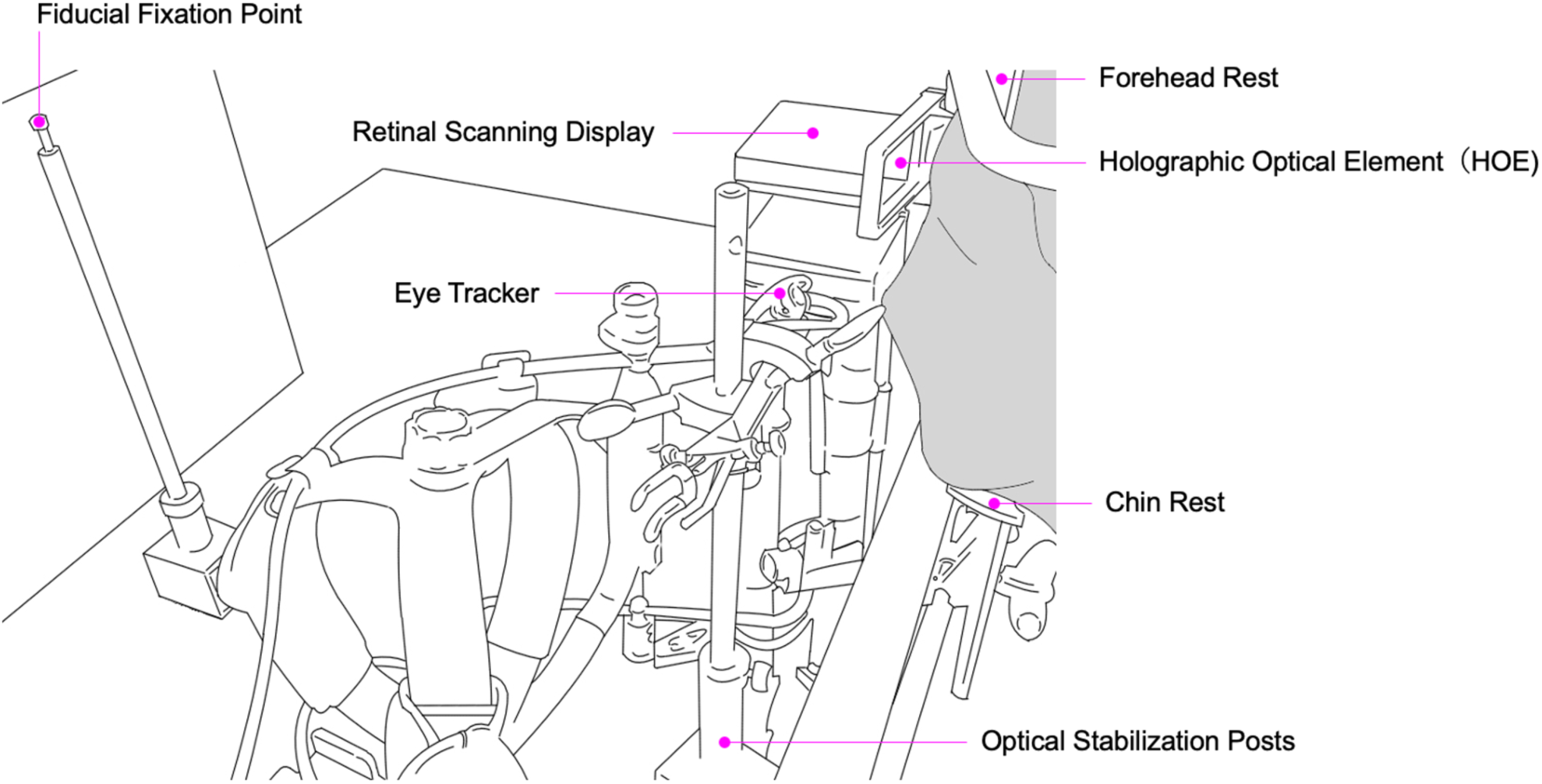
Schematic of experimental setup and visual stimulation

## STIMULATION

Figure 5 is a schematic of visual stimulation as it enters the eye and falls on the retina. We generated graphics for the participant task using a Windows 10 PC running Psychophysics Toolbox (D. H. Brainard, 1997; Kleiner et al., 2007; Pelli, 1997) for Matlab (Mathworks, Natick, USA).

**Figure 5.**
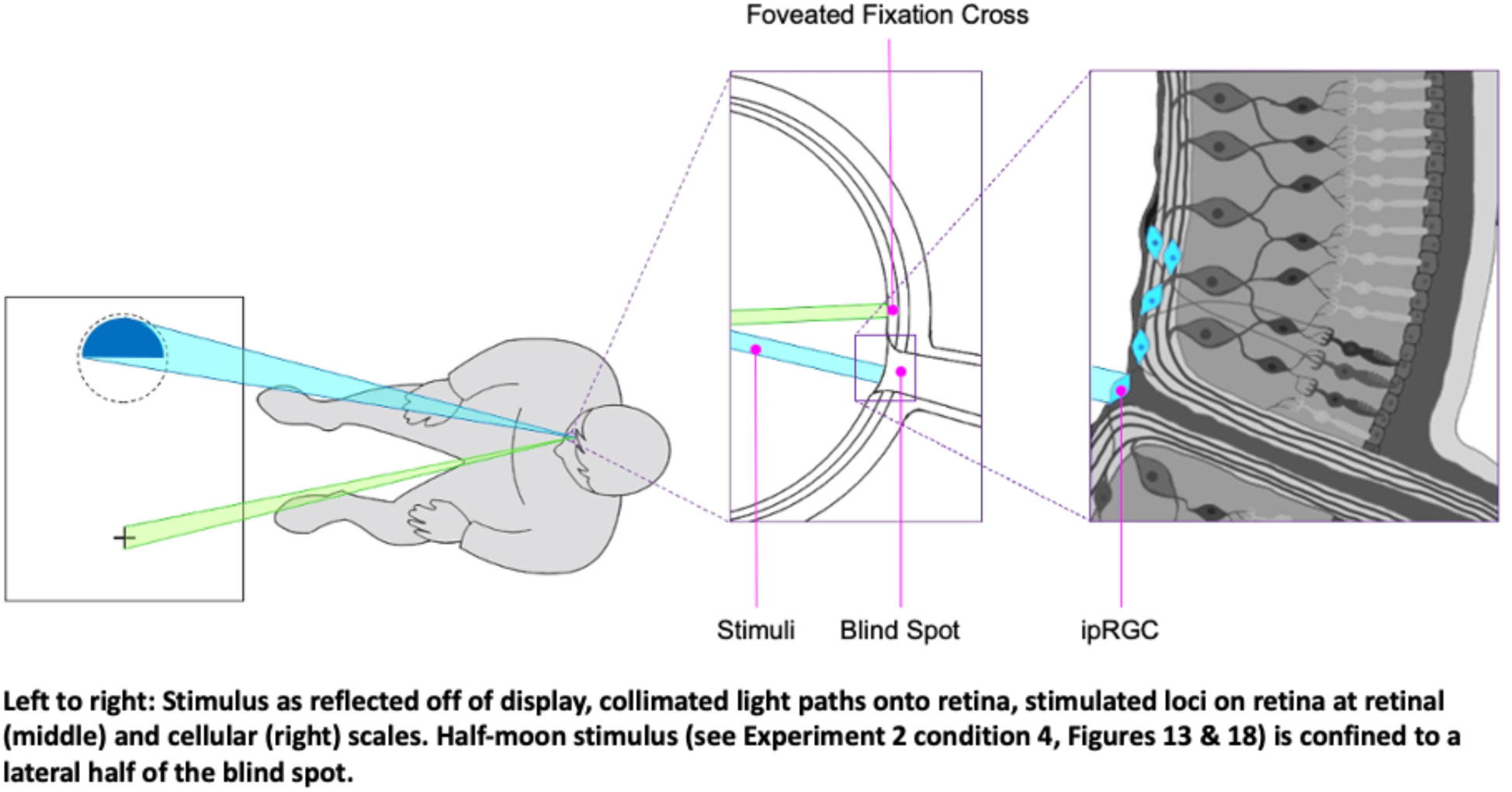
Schematic of Spatially Confined Stimulus Delivery, Across Scales, With Respect to the Targeted Region of the Retina

**Figure 6.**
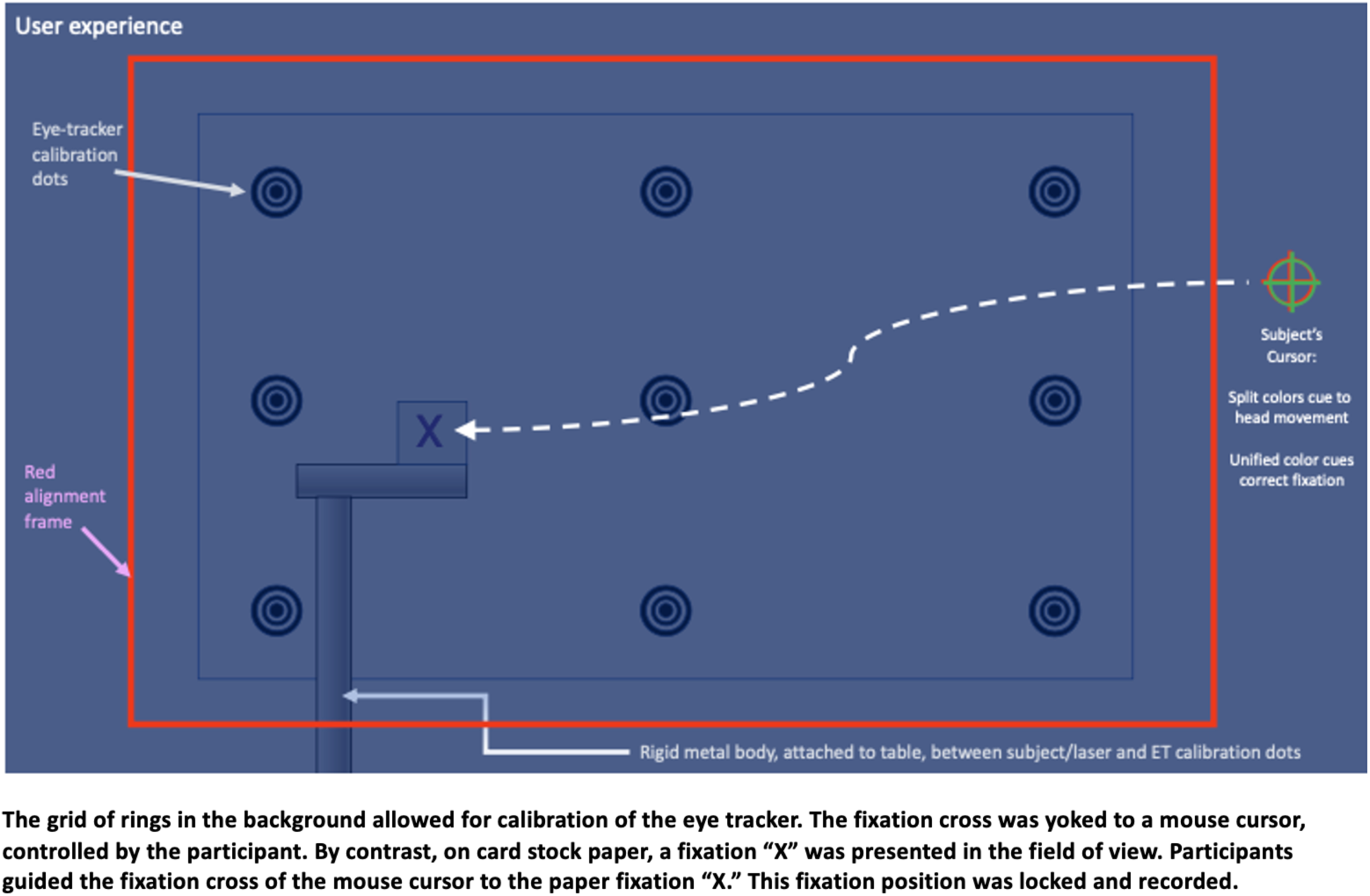
Schematic of the Field of View From a Participant’s perspective

**Figure 7.**
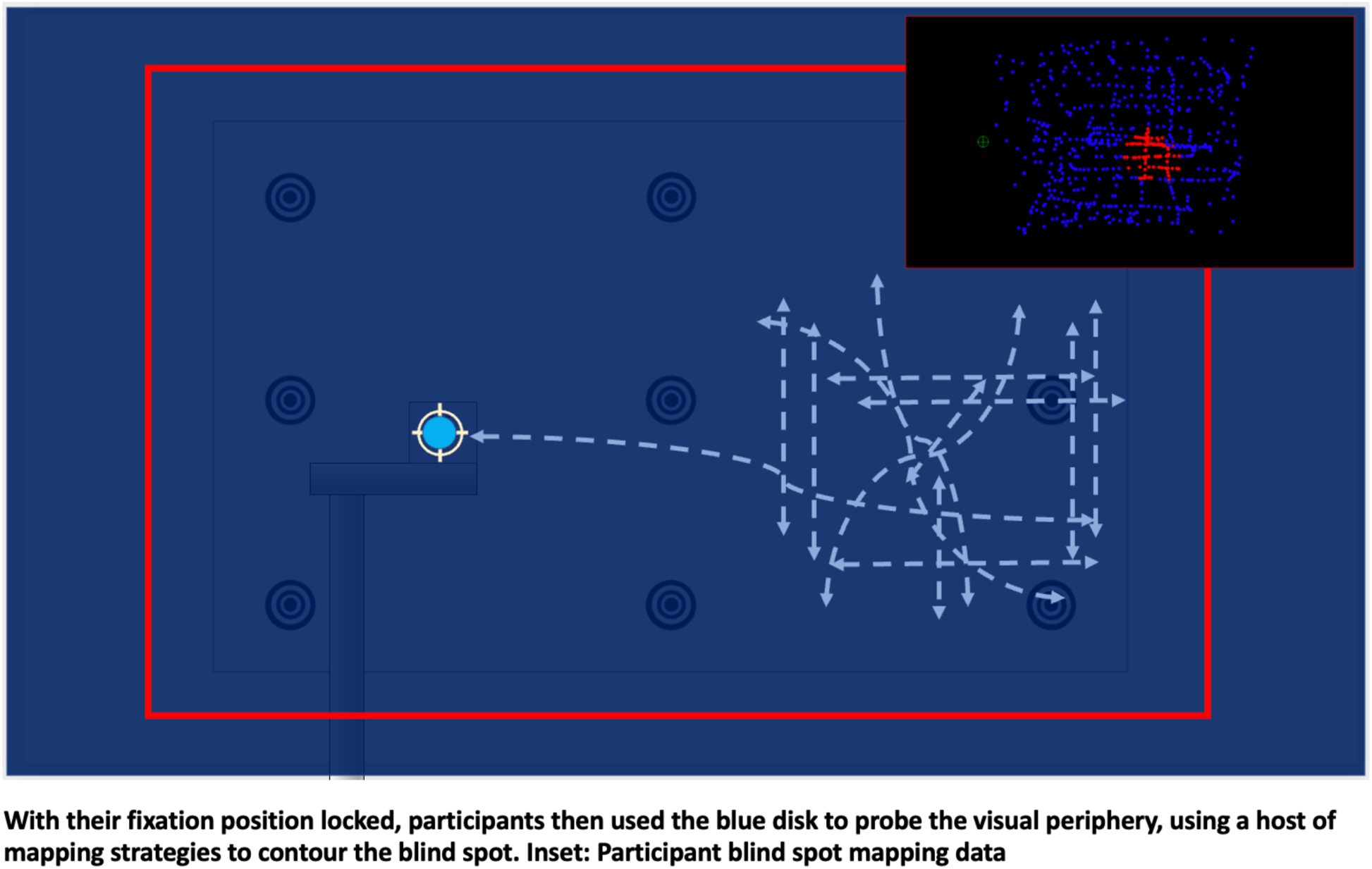
Blind Spot Mapping

Blind Spot Mapping and Optically-Constrained Gaze Fixation

Every experiment started by mapping the participant’s blind spot. Based on the established wavelength preference of melanopsin, we used blue light stimuli at a wavelength of 445 nm, below the established preference of melanopsin (460–500 nm). Head-stabilized participants were guided through the process of using the RSD to map their own blind spot, while maintaining constant gaze-fixation on a gaze-contingent fixation cross. The optical characteristics of the RSD limited the conditions of viewing; Were a participant to saccade off of the fixation cross, the entire field of visual stimuli would disappear. Further, participants received feedback about small head movements based on the appearance of the fixation cross. If the head moved out of place, the yellow fixation cross would split into its red and green components due to differential diffraction off of the display element.

Having established fixation, they used a computer mouse that controlled a blue target, blinking at 0.5 Hz. They maintained gaze-fixation, while also left clicking as they moved their mouse cursor toward the right visual periphery. Wherever they could no longer see the blinking target, they would right click. When the blinking target would reappear, they would again left click. Using a variety of movement patterns, gaze-fixed participants mapped their peripheral visual field with respect to their sensitivity to a flashing blue disk. The diversity of blind spot sizes, varying significantly in their outer contours, highlighted the importance of this subject-specific approach. Following the mapping procedure, stimuli could be presented with respect to the contours of the blind spot. During the experiments, we measured pupillary responses, enabling subsequent analyses often used to infer the integrity of visual pathways.

## 6. Experiments

### Participants and IRB Approval

Across all experiments, we ran a total of thirty, healthy individuals of mixed age range (19-66). They had normal or corrected to normal vision, with glasses removed for the experiment (the laser image remains in focus regardless of acuity or lens accommodation). None of the participants had neurological or visual disorders. All experiments and their procedures were approved by the Committee for the Protection of Human Subjects at the California Institute of Technology (Protocol numbers 21-1157 and 21-1161) and the Sony Bioethics Committee (Protocol number 21-F-0031), respectively. All participants gave written informed consent prior to the study.

### Experiment 1 The initial exploratory observations

#### Participants

During experiment 1, ten participants were recruited from within our lab due to task-demands for maintaining stable fixation (Age: 19-66).

#### Procedure

We conducted an exploratory experiment using the setup and the blind spot mapping procedure that we developed. Participants did not take time for dark adaption before starting the experiment. Following blind spot mapping, we did not expect dark adaptation to be essential to the detection of immediate pupillary responses to illumination. Participants performed our blind spot mapping procedure, immediately followed with the presentation of a blue disk, blinking at 0.5 Hz within the participant’s blind spot for one minute on, and one minute off.

Disk size varied so as to fit well within the participant’s blind spot. The blinking disk was presented for one minute, then turned off for one minute, in alternation. Timeseries plots of pupil size were examined visually. Windowed averages for the stimulus on and off periods were computed.

In this exploratory phase, we employed electroencephalography in order to examine neural responses to blue stimuli. Rhythmic visual stimuli have been demonstrated to induce measurable changes in electroencephalography (EEG; Moratti et al., 2007; Vialatte et al., 2010). The frequency spectrum of these electrophysiological responses are known as steady-state visual evoked potentials (SSVEP), and have been thoroughly investigated for decades (Moratti et al., 2007; D. Regan, 1978; M. P. Regan & Regan, 1988; Vialatte et al., 2010). We presented a small, blinking blue disk in the blind spot at test frequencies of 0.5 Hz, 5 Hz, or 7 Hz. EEG was recorded continuously from 64 electrodes using a Biosemi ActiveTwo system, digitized at a rate of 2048 Hz and downsampled to 250 Hz (Moratti et al., 2007). In analysis (Figure 9), we explored multiple different settings, filtering signals between 0.1 Hz and 40 Hz, as well as between 0.1 Hz and 20 Hz. Multiple reference channels were tested, including Cz and FPz. The frontal channel was tried, as it can enhance the signal from the posterior visual cortices. Figure 9 data are referenced to Cz.

## Results

A key development in experiment 1 was the execution of a personalized protocol to map an individual’s blind spot. The results of this mapping procedure revealed variability in both the shape and size of individuals’ blind spots (Figure 8). By contrasting four participants’ maps, in figure 8, it is evident that participants were able to employ a variety of strategies to mark the visibility and invisibility of the blue disk.

**Figure 8.**
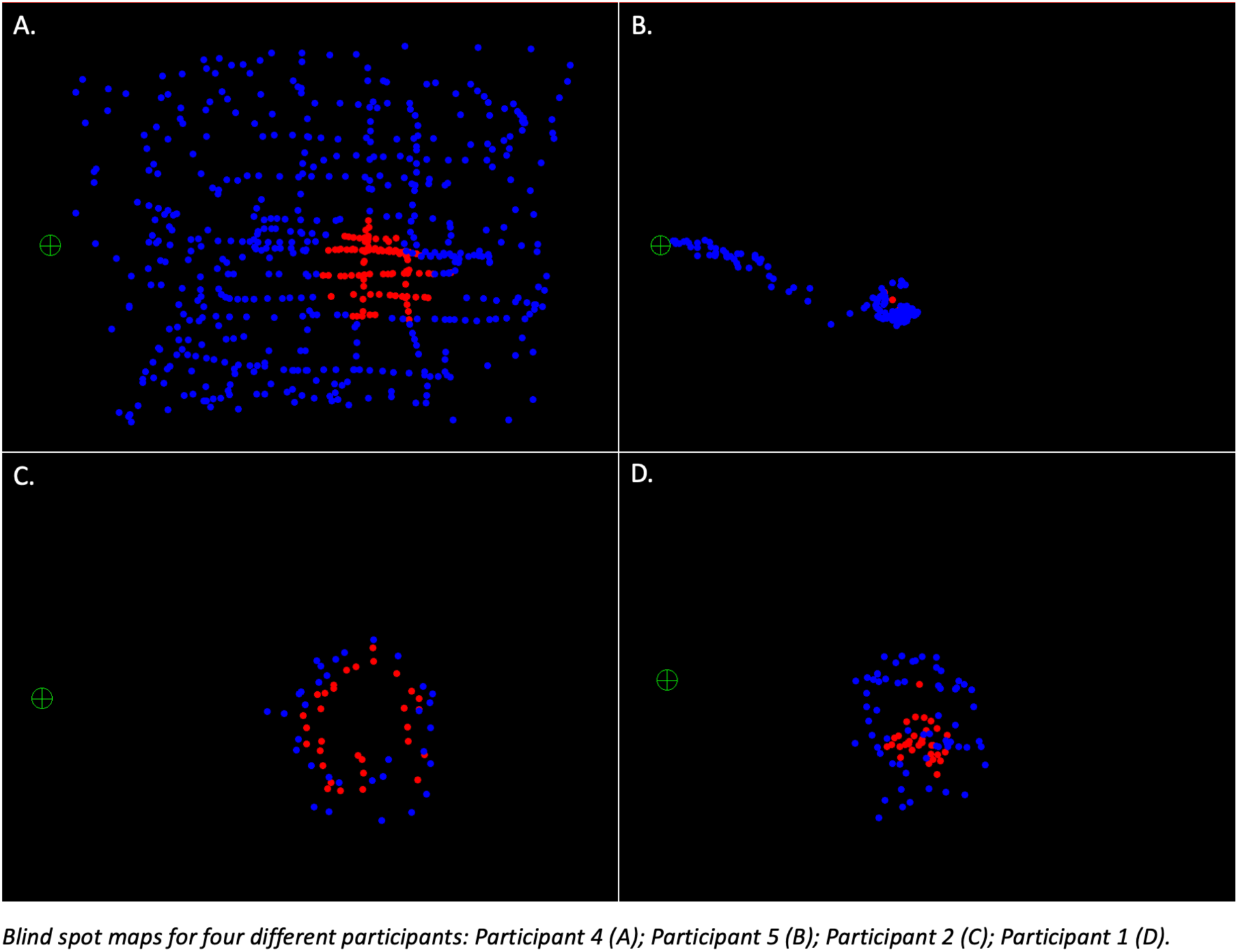
Blind spot maps for four different participants

**Figure 9.**
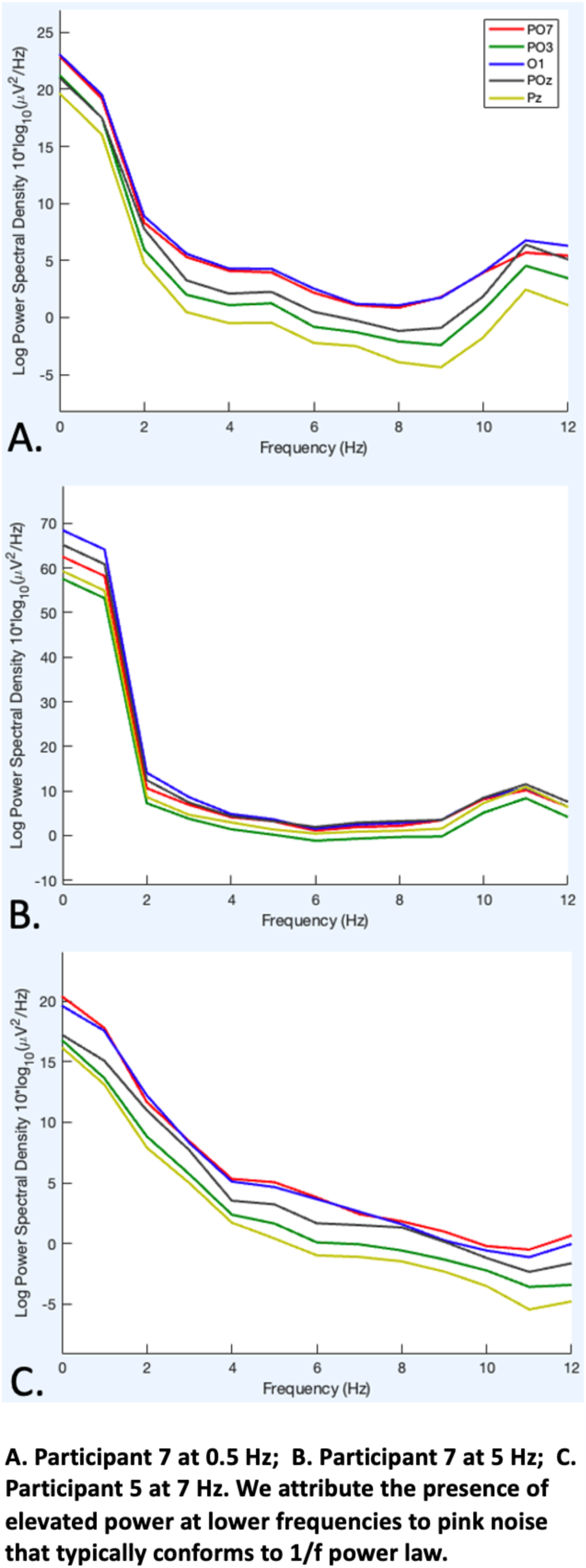
Frequency Spectra for Five EEG Channels Over the Occipital Lobe

While data from this exploratory experiment was highly noisy, it did allow tuning of the device and experiment. Additionally, we identified an alternative measure of pupillary response, which we discuss in experiment 2, that is tuned to melanopsin-associated activity. Alongside measures of pupil size and EEG from a subset of participants, participants’ phenomenological reports, such as visual artifacts, were systematically logged.

Our EEG analysis did not show signs of an SSVEP in response to rhythmic presentation of a small blue stimulus (445 nm) to the blind spot. Lee et al. (2018) stimulated the peripheral visual field at frequencies partly overlapping with our own, finding that the oscillatory response, of specific peak enhancement at the stimulus frequency within the Fourier spectrum, could be decoded from EEG data (Lee et al., 2018). Yet, our presentations to the blind spot did not yield visibly identifiable oscillations (i.e., enhancement of power in spectrum plots) at our test frequencies of 0.5 Hz, 5 Hz, or 7 Hz (Figure 9). This failure could be due to any of several reasons. It is possible that, while ipRGCs would respond by sending out sufficient periodic signals, the visual cortex lacks physiological sensitivity to our stimuli at the blind spot. Another possibility is that our stimulation was too weak to significantly activate ipRGCs. It could also be the case that SSVEPs were present within cortex, but we failed to record them for technical reasons. While impedances were monitored, it is possible that we failed to control for both motor-related and electrical sources of noise. Since participants’ cheeks and forehead frontally rested against a stabilizing assembly, and abutted the RSD to the front and right side, ocular activity could not be recorded through placement of electrodes on the face. Indeed, anterior channels were also saturated with electrophysiological noise due to pressure from the headrest. It is also possible that a stimulus-driven oscillation may be hidden by the prominence of the typical, low frequency noise that surrounds our flicker frequency. Future experiments might need to increase the size or intensity of stimulation, or better detect the presence of smaller oscillations by removing aperiodic components of the frequency spectrum, such as slope.

During experiment 1, across participants, we discovered two common visual artifacts. The first artifact was a duplicate image of the target that was approximately five degrees to the right of, slightly lower in luminance than, and time-locked with the target, or the “ghost” artifact. The other artifact was a diffuse cloud of light typically reported when the target was within, but near the interior boundaries of, the blind spot, or the “halo” artifact. For example, this was the case when the target disk’s diameter was significantly increased without exceeding participants’ visual detection threshold. So far, the underlying causes of these artifacts remain technically unsolved.

To minimize recurrence of artifacts, we developed the practice of reminding participants to align their heads with the right-side plane most reliably resistant to artifact. The halo artifact could be significantly mitigated by maximizing the distance of the invisible, or blind spot-occluded target from the edges of the blind spot. Notably, however, this is necessarily in tension with increasing the luminance energy presented to ipRGCs within the blind spot. The issue of the ghost artifact, however, could not be resolved, and was a source of confusion for participants unsure of the true identity of stimuli emerging from the right bank of the blind spot. That is, they were unsure whether the emerging disk was the true target or the right-leading ghost-target. The causes of these artifacts may be explained by crosstalk between incoming channels of light, due to differences in the grating periods of red, green, and blue light, diffracted across the planar diffraction gratings and hologram layers of the HOE (Collier & Pennington, 1967; Mukawa et al., 2009). These artifacts may have contributed to the noise within, and non-significance of, our obtained data.

### Experiment 2 The examination of pupillary responses

#### Procedure and Stimuli

In the second experiment, we optimized the methods developed in experiment 1 and deployed four stimulus conditions. Stimuli were presented continuously for 30 seconds, followed by 60 seconds off, repeating 4 cycles. In condition 1, the control condition, a disk was presented outside the blind spot and visible to the participant (Figure 10). In condition 2, the first experimental condition, the disk was shown inside the blind spot (Figure 11). In condition 3, we examined whether activation of melanopsin in the blind spot would have a modulatory effect on the pupillary response to visible stimuli. The stimuli consisted of the same disk inside the blind spot, plus a ring which extended across the boundary of the blind spot (Figure 12). Such rings are known to be perceptually “filled-in.” Because the hole in the ring is entirely within the blind spot, the ring is seen as a solid disk. Would presentation of an actual disk inside this hole result in modulation of the pupillary response, via activation of melanopsin? Modulation of both short- and long-latency pupillary responses was examined. In condition 4, we tested for interactions between rod and cone inputs with melanopsin inputs as they are combined within individual ipRGCs. An ipRGC receives rod and cone signals from a constrained region of the visual field, and transmits that signal down an axon as it exits the eye through the corresponding location in the optic disk. Thus, activity from stimulation of the right side of the fixation cross and visual field will be routed through axons running through the right side of the blind spot. To see if the two signals interact, we illuminated the right side of the visual field in green, leaving a hole in the blind spot. We then showed either a blue or a red “half-moon” stimulus on the right side of the blind spot (Figure 13).

**Figure 10.**
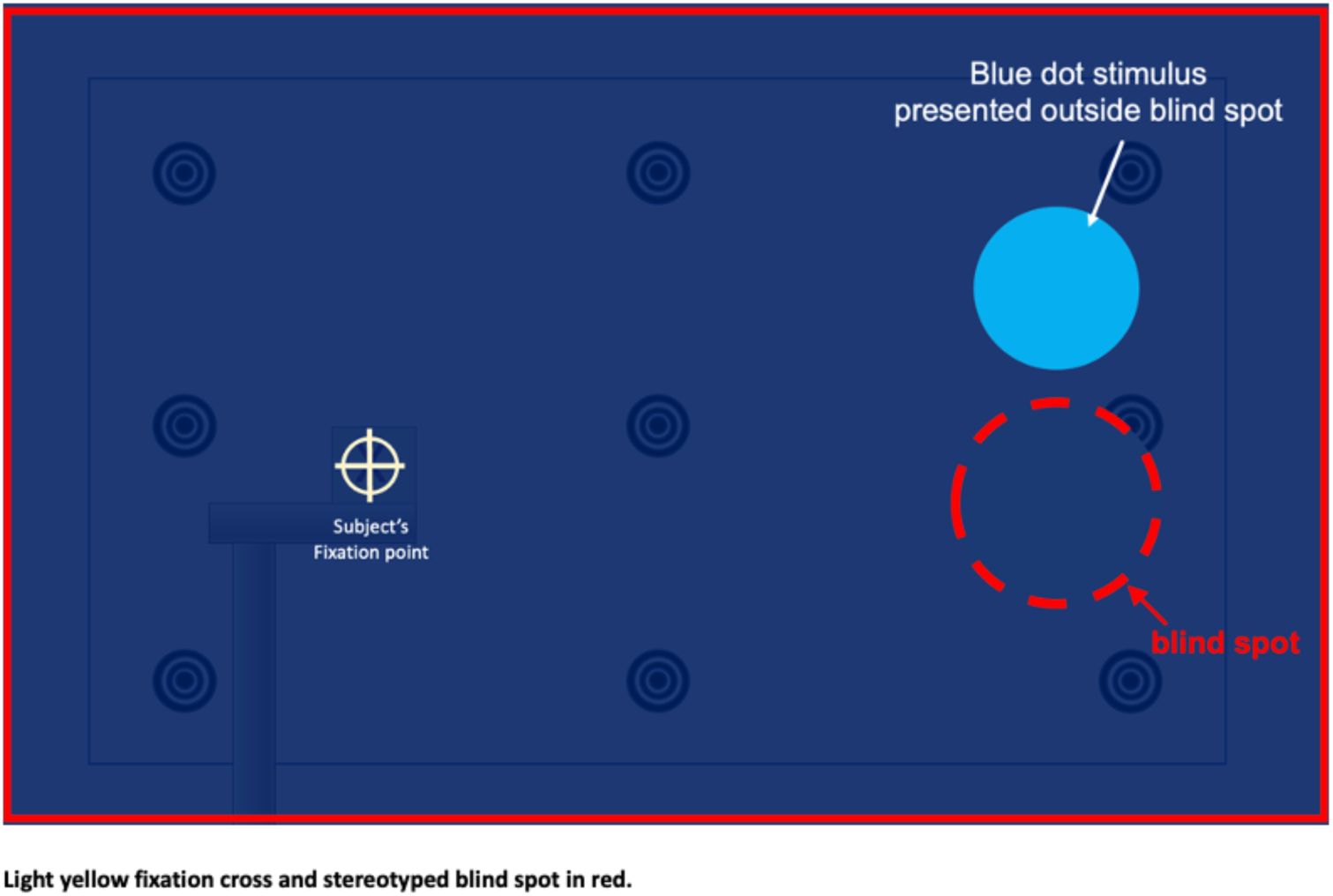
Schematic Diagram of Stimulus Condition 1, Baseline Stimulation Outside of Blind Spot

**Figure 11.**
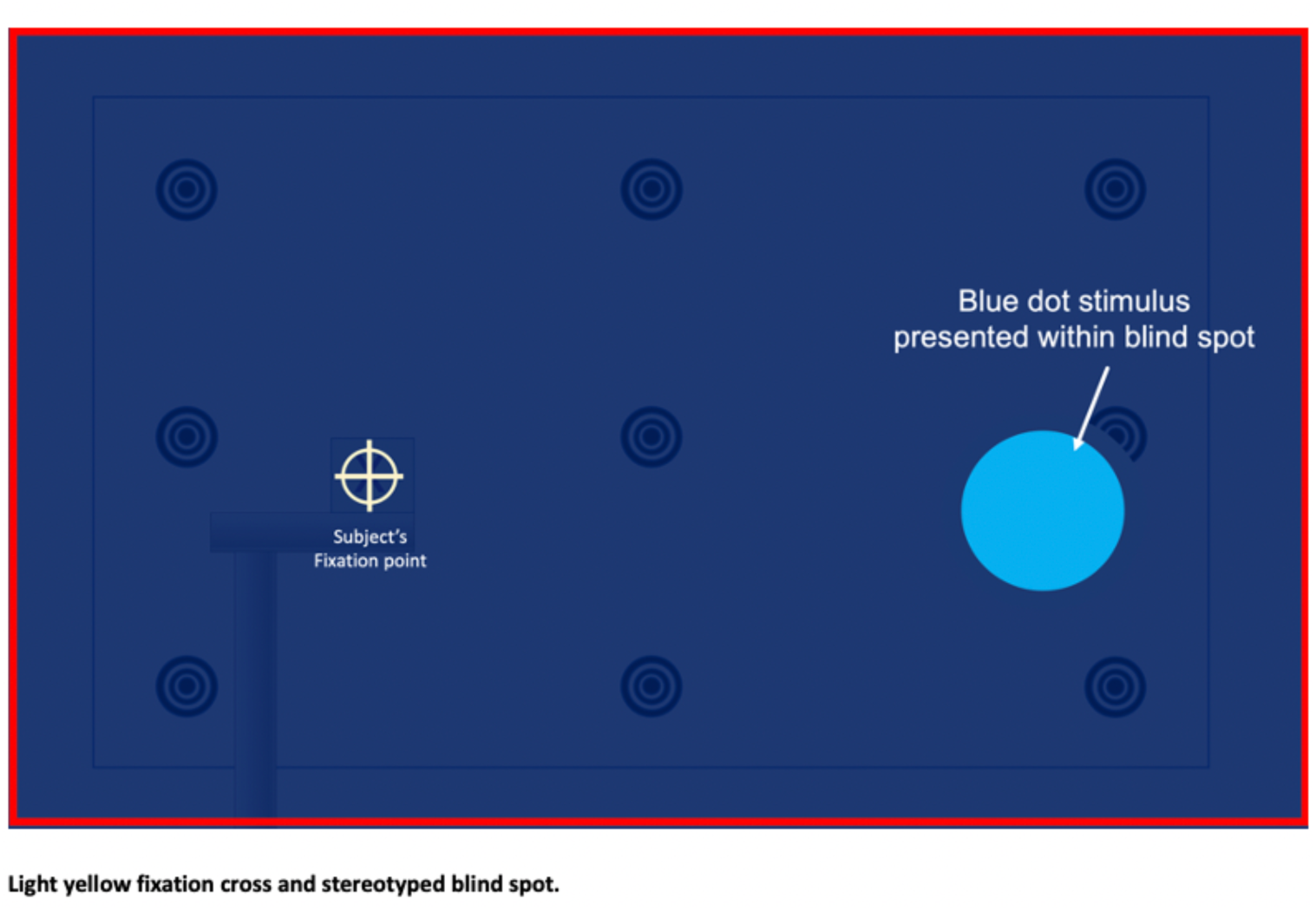
Schematic Diagram of Stimulus Condition 2, Stimulation Inside of Blind Spot

**Figure 12.**
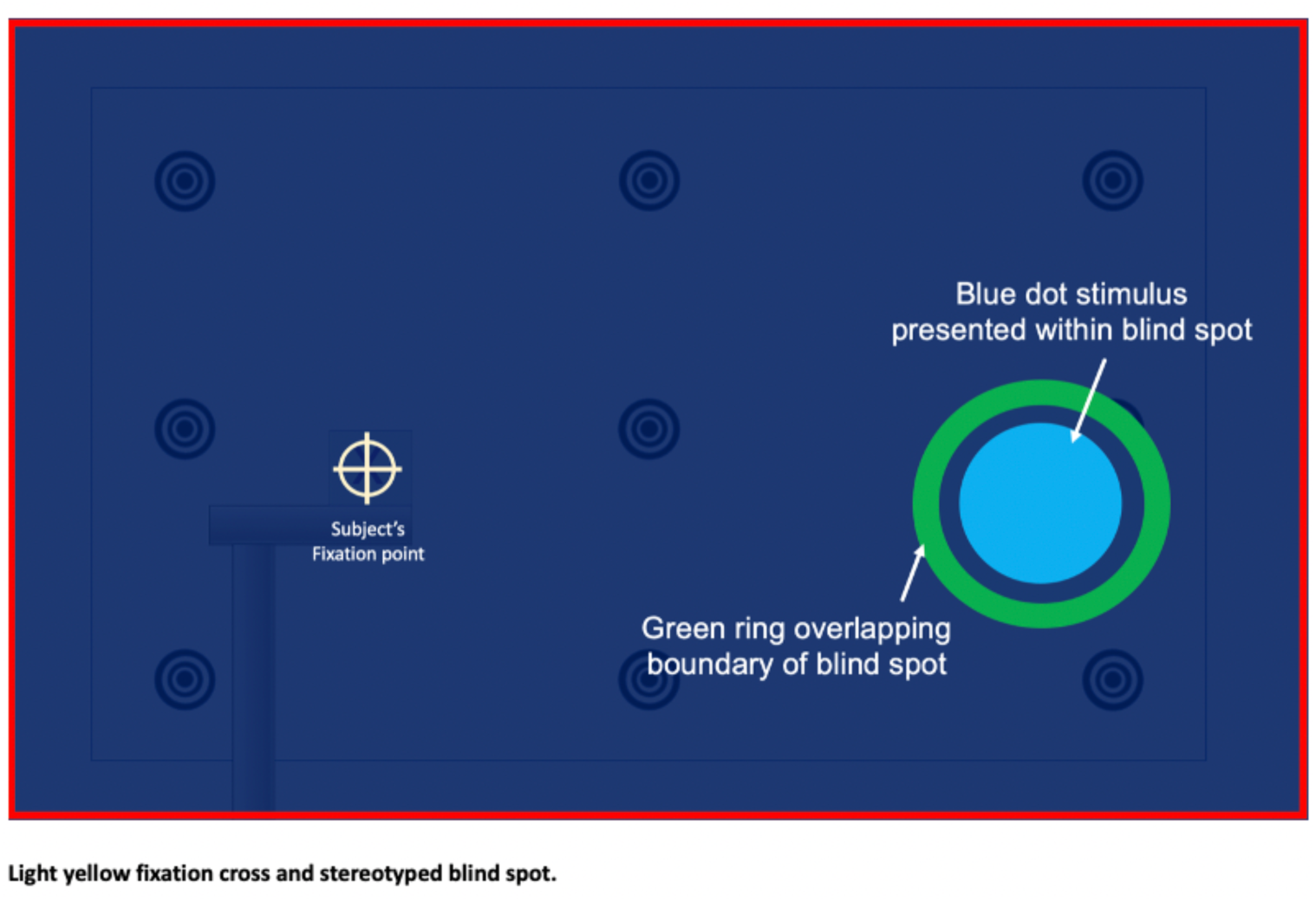
Schematic Diagram of Stimulus Condition 3, Green-ring Across the Boundary of a Stereotyped Blind Spot

**Figure 13.**
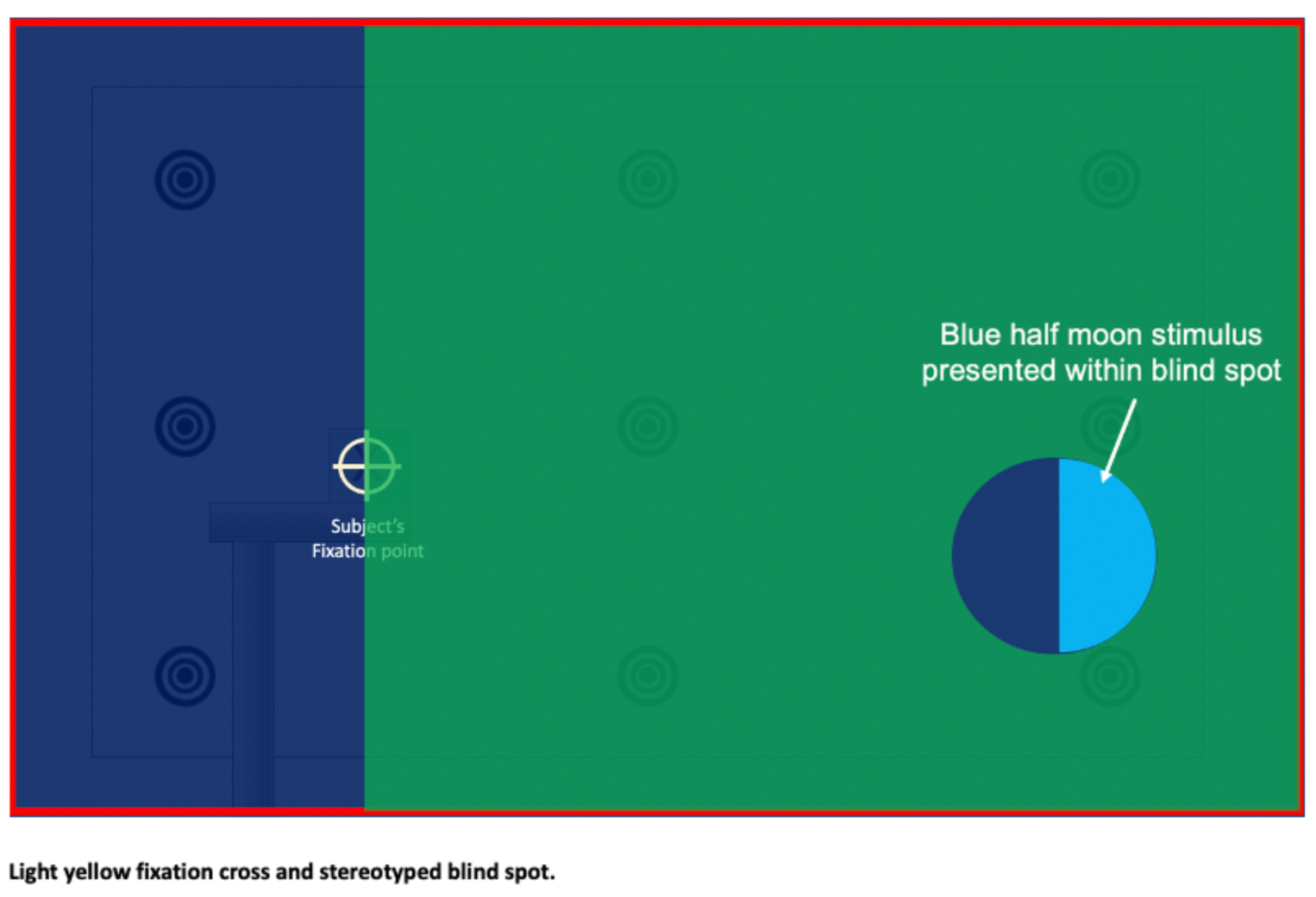
Schematic Diagram of Stimulus Condition 4, Half-disk Stimulus Inside Blind Spot, With Green Background Filling the Right Visual Field

These conditions tested for implicit responses to stimulation within the blind spot, measuring both the pupillary light response (PLR) and the post-illumination pupillary response (PIPR). Whereas the PLR occurs rapidly in response to stimulus onset, the PIPR occurs over a prolonged period after the stimulus is turned off. The PLR is typically used to measure both inner and outer retinal function, and the PIPR is used specifically to measure the function of ipRGCs (Adhikari et al., 2015). The stimuli and analyses were based on monkey and human experiments (Gamlin et al., 2007). PLR was calculated as the percent change in pupil size from baseline (10 sec pre-stimulus average) to stimulation period (average for time window of 10-18 sec after onset). PIPR was calculated as percent change in pupil size from baseline (10 sec pre-stimulus average) to post-stimulus period (average for time window of 15-30 sec after offset).

#### Participants

During experiment 2, thirty participants were recruited in total. They ran overlapping subsets of the conditions. They were naïve as to the experimental conditions and hypotheses and did not participate in previous experiments. However, the demands of the experiment turned out to require experienced psychophysics participants who could perform the tasks with the stability required. Specifically, five expert participants participated in all four conditions (Age: 19-66). The pupil data presented are based on five expert participants who were run in all conditions. The number of participants remains too low for statistical tests. For this reason, we present results with descriptive statistics to describe patterns in the data. These statistics can be used for future experiments to perform power analyses and to determine appropriate sample and recruitment sizes.

### Results

#### Condition 1

Figure 14A shows the PLR when the disk, outside and above the blind spot, was blue vs. when it was “black,” with black equating to no stimulus, given a black background—for which the display image remains blank and transparent. When the disk was blue, on average, the pupil constricted by around 5% (mean size=94.99%, SEM=3.28%, N=5). When the disk was “black” (i.e., no stimulus), the pupil showed little change (mean=99.19%, SEM=1.4%, N=5), aligning with expectations and literature.

**Figure 14.**
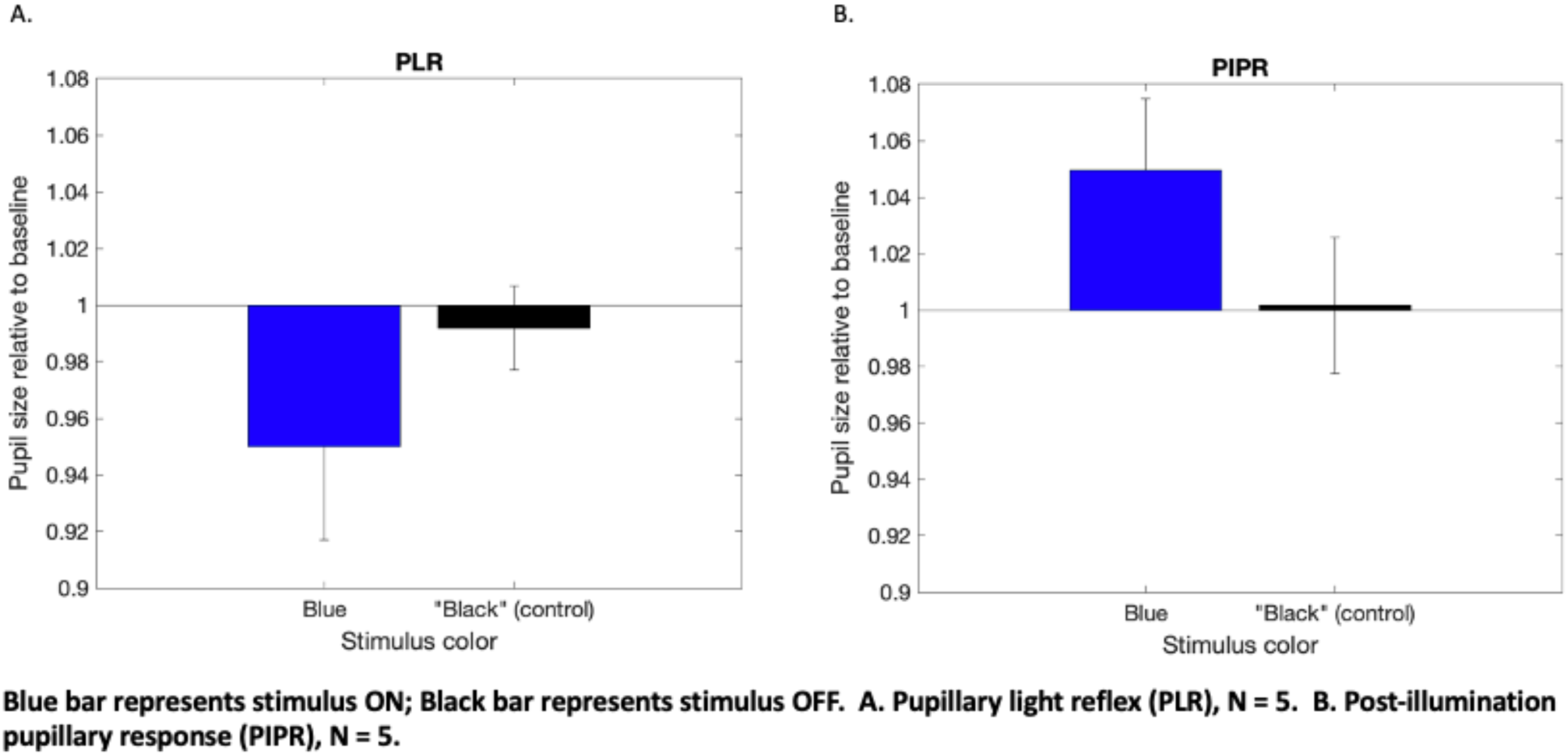
Presentation Outside and Above the Blind Spot

#### Condition 2

Figure 15A shows the PLR when the disk was blue vs. when it was red (643nm, outside the range of melanopsin activity). When the disk was blue, the pupil again constricted around 5% (mean=95.3%, SEM=2.89%, N=5) When the disk was red, the pupil showed little change (mean=100.5%, SEM=2.18%, N=5). This is consistent with expectations, where melanopsin would respond to blue but not to red light.

**Figure 15.**
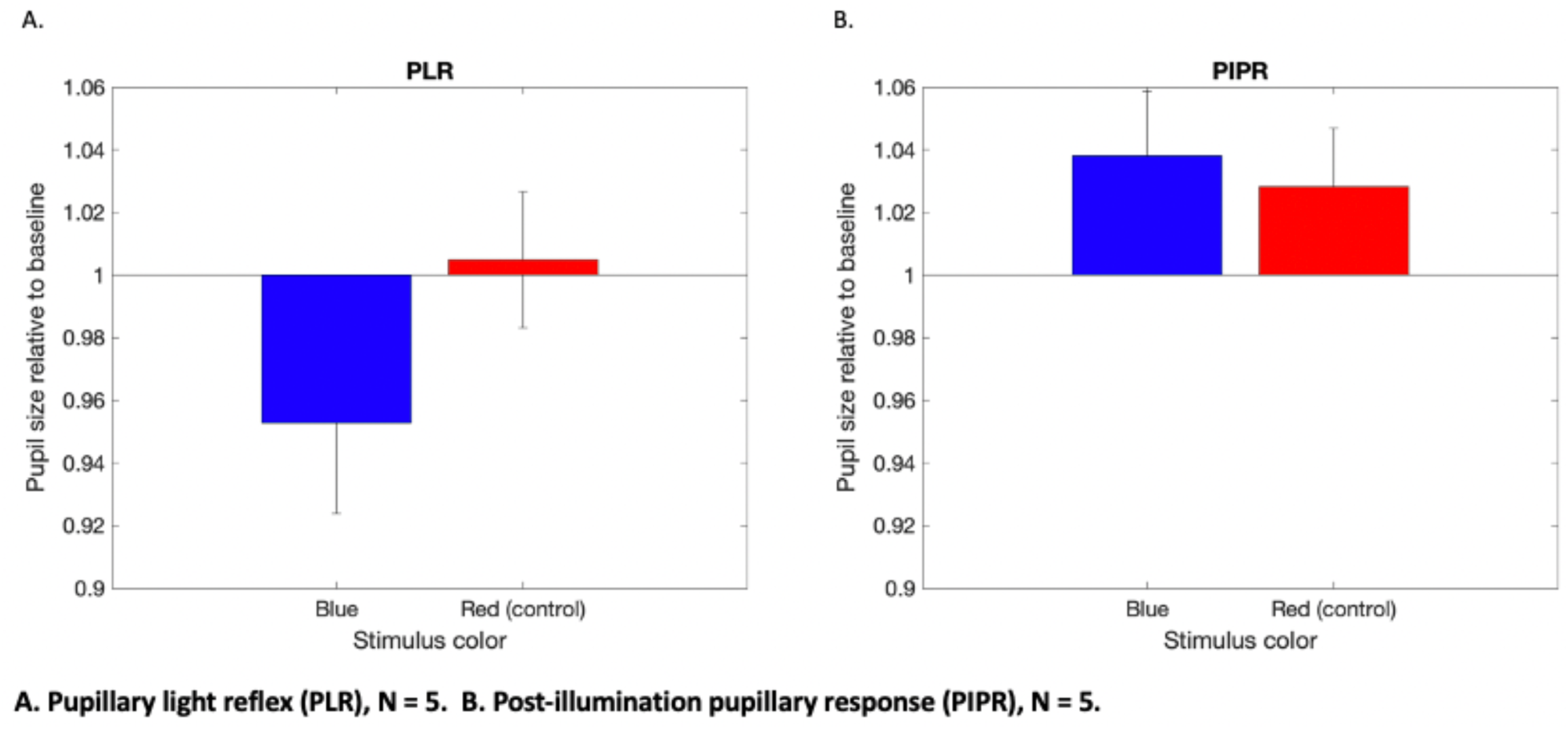
Stimulus Presentation Inside the Blind Spot

**Figure 16.**
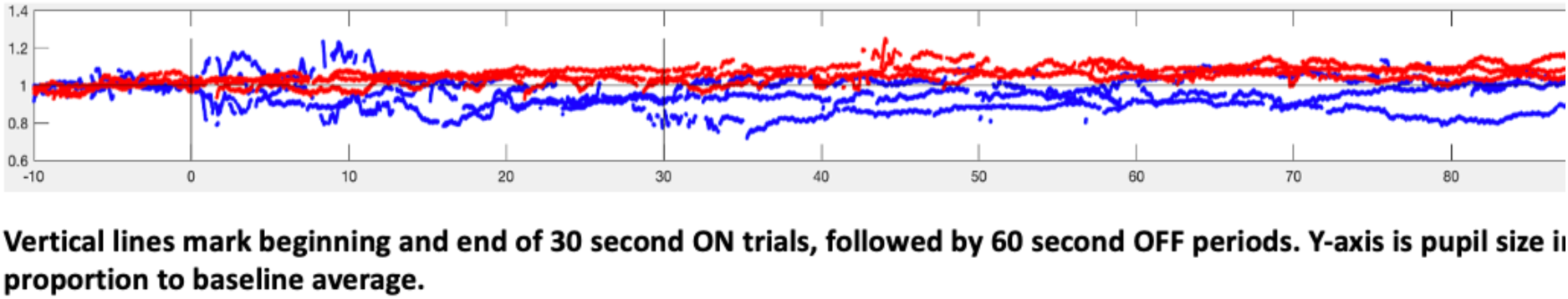
Full time course of PIPR, in seconds, for one participant and reflecting color of stimulus.

Analogous results can be seen, in these same conditions, when looking at the PIPR to the visible control location in condition 1 (Figure 14). Figure 14B shows the PIPR in response to the disk shown within the blind spot, blue vs. “black.” After the offset of a blue disk, the pupil dilated, on average, 4.95% beyond the baseline size (mean=104.95%, SEM=2.55%, N=5). After a “black disk” (i.e., no stimulus) there was little change (mean 100.17%, SEM=2.42%, N=5).

For presentations within the blind spot (condition 2), the PIPR was similar for the red (643nm) and blue (445nm) stimuli (Figure 15B). After offset of the blue disk, the pupil dilated on average 3.81% (mean=103.81%, SEM=2.05%, N=5). These measurements after the red stimulus were initially surprising. After offset, the pupil dilated, on average, 2.84% (mean=102,84%, SEM 1.86%, N=5). We had not expected red stimuli to have any effect inside the blind spot. Even more mysterious, we were seeing a post-stimulus dilation, even though the pupil had not constricted in response to the red stimulus turning on.

Upon closer examination of the raw data, however, we discovered that this dilation was not actually due to the red stimulus, but in fact was the continued dilation from the offset of the blue stimulus from the previous trial. Figure 12 shows the continuous traces of pupil size through a series of trials. Red and Blue stimuli alternate in this experiment, and their traces are overlaid. Blue traces reflect the pattern described above, where there is pupil constriction during stimulation, and dilation after stimulus offset. The trace continues to rise even 60 seconds after the stimulus offset, when the next (red) stimulus is set to begin. The red traces, meanwhile, continually rise through the whole trial. Lining up the traces end-to-end, it is apparent that the rising red traces are just the continuation of the rise still occurring at the end of the blue traces. Looking back through Gamlin (2007), on which we based our design, they indicated that participants maintained fixation for 16-30 seconds after stimulus offset, but did not explicitly say when the next trial would start. It would seem that their duration of required recovery must have exceeded the duration of intertrial interval that we used. This is suggested by other literature, wherein extremely prolonged recovery periods were used, with some studies leaving as much as 7 minutes of inter-trial interval to allow for recovery (Joyce et al., 2015).

#### Condition 3

Here, we found that the green ring caused much higher pupillary responses across the board, with high variances. Figure 13 shows the PLR (17A) and PIPR (17B) to these ring stimuli. Green stimuli with blue inside caused a PLR constriction of nearly 10% (mean=90.18%, SEM=6.05%, N=5), and green stimuli with red inside more than 16% (mean=83.15%, SEM=4.82%, N=5). For PIPR, blue stimuli caused a dilation of nearly 15% (mean=114.29%, SEM=4.40%, N=5) and red stimuli over 16% (mean=116.37%, SEM=4.84%, N=5). Numerically, the responses in the red-centered green rings are larger, but this is hard to interpret due to the high statistical variance values.

It may be that the green component of the stimuli may have been too strong in these conditions. Its contribution may have been too dominant, leading to intense pupillary responses that left no room for further modulation by the blue and red blind spot components. Both the high values and high variances in pupil response suggest that the blue and red components could be washed out. Future experiments could test lower intensities of green light, which may require hardware modification of the green laser channel.

#### Condition 4

Based on Miyamoto and Murakami (2015), we expected that a blue half-moon stimulus on the right side of the blind spot would modulate the pupil response to the green stimulation of the right visual field. Unfortunately, we ran into similar issues as with the green rings in the previous section. With the green field illumination constantly on, the addition of blue or red moons had minimal effect. Figure 18 shows the PLR and PIPR measurements, respectively. PLR was minimally affected by blue half-moons (mean=99.89%, SEM=2.15%, N=5), and the same for red half-moons (mean=100.28%, SEM=2.78%, N=5). Similarly, PIPR effects were minimal for blue (mean=102.94, SEM=3.73%, N=5) and red (mean=98.27%, SEM=3.06%, N=5). Here too, it appears that the green component of the stimuli should be tested at a lower level.

**Figure 17.**
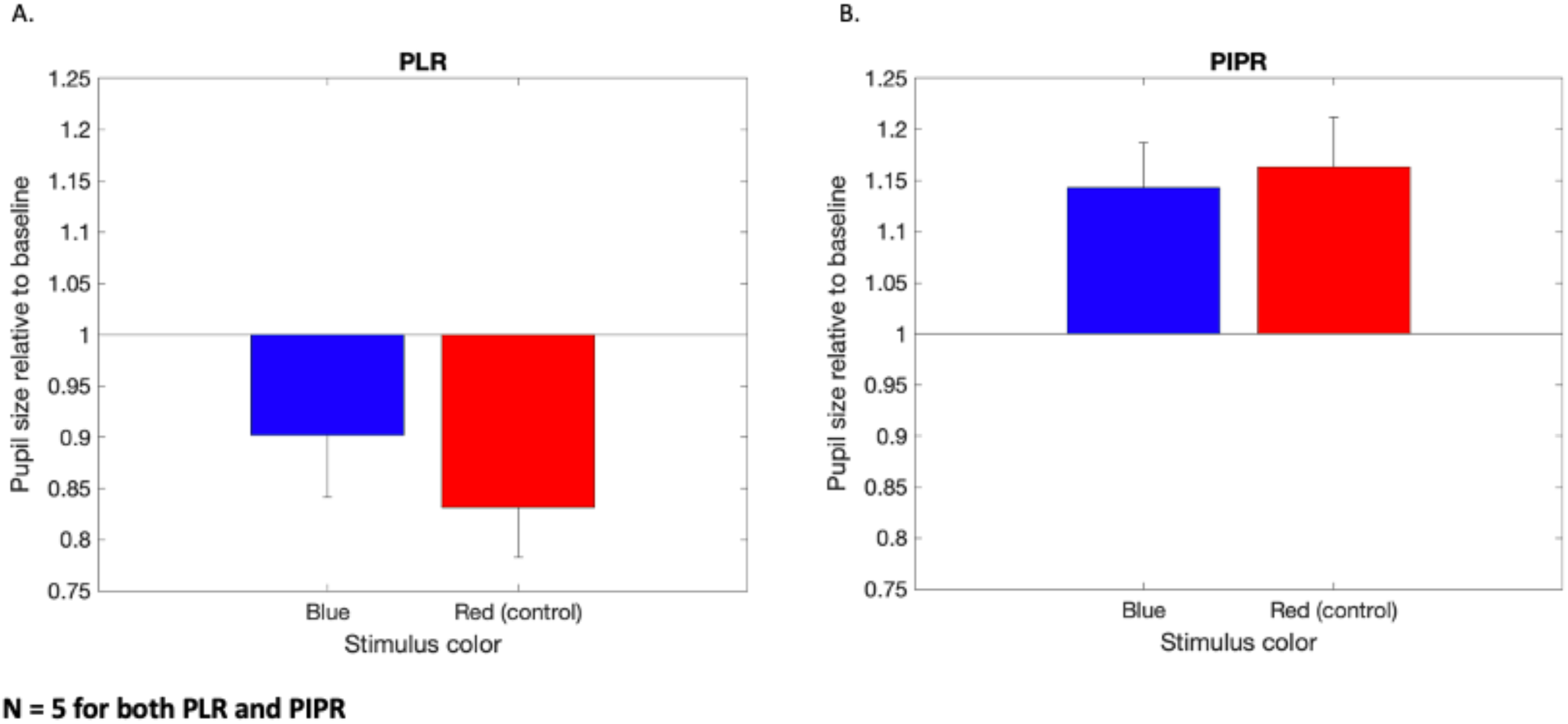
Stimulus Presentation Inside the Blind Spot, With a Green Ring Across the Boundary of the Blind

**Figure 18.**
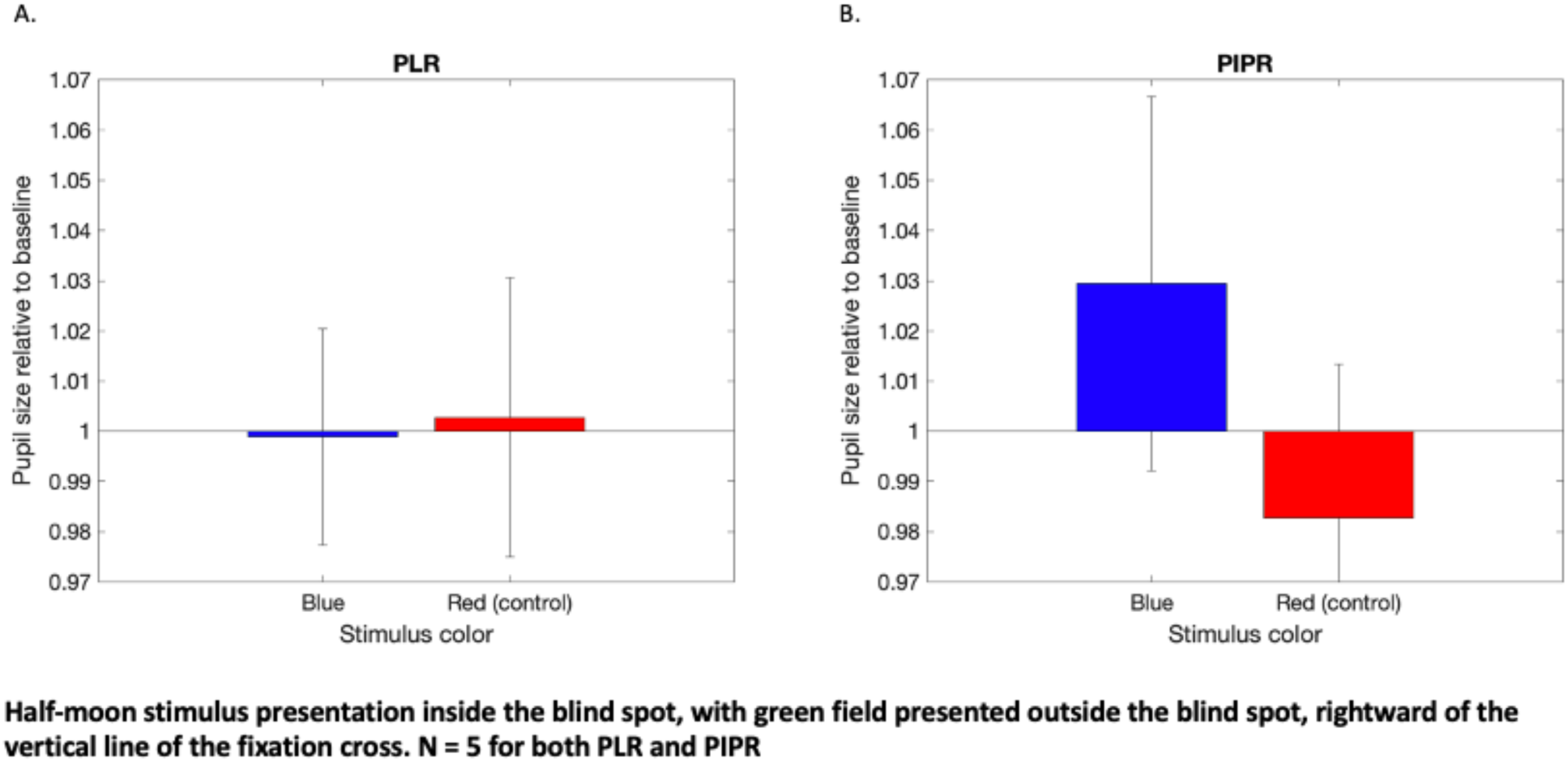
Half-moon stimulus presentation inside the blind spot, with greenfield outside

### Experiment 3 The inference of implicit visual perception arising from the blind spot

Following the second experiment of data collection and analysis, we aimed to identify a role of ipRGCs in implicit visual processing. It is known that the eye continually moves even when a person attempts to hold a target in their gaze. These deflections have been found to be biased if the person’s attention is directed covertly in some direction. Since participants’ gaze was always fixed on the fixation cross, we hypothesized that the average direction of deflection away from the cross might indicate a participant’s implicit sensitivity to the presence of a stimulus that was invisible to their explicit awareness. If so, we might expect that eye movements would be biased in the direction of the blind spot when a stimulus was displayed there.

#### Participants

During experiment 3, gaze position data from 10 participants from experiment 2, condition 2, were analyzed (Age: 19-66). They were naïve as to the experimental conditions and hypotheses, and did not participate in the previous experiments.

#### Procedure

Data were drawn from experiment 2, condition 3, throughout which participant were instructed to maintain their gaze on the fixation cross. Gaze position was plotted for stimulus ON vs stimulus OFF periods. To remove eccentric jumps in gaze position, we omitted data beyond a threshold of three standard deviations away from the fixation position.

### Results

Figure 19 highlights data from a sample observer. The fixation point is indicated by the red cross. For the ten participants, the two columns on the left show the sampled gaze positions during stimulus ON and OFF, respectively. Each of these two columns indicates the relative position of the blind spot to the right of fixation. The column on the right indicates the average direction of off-fixation gaze during both stimulus ON (dark blue) and stimulus OFF (light blue).

**Figure 19.**
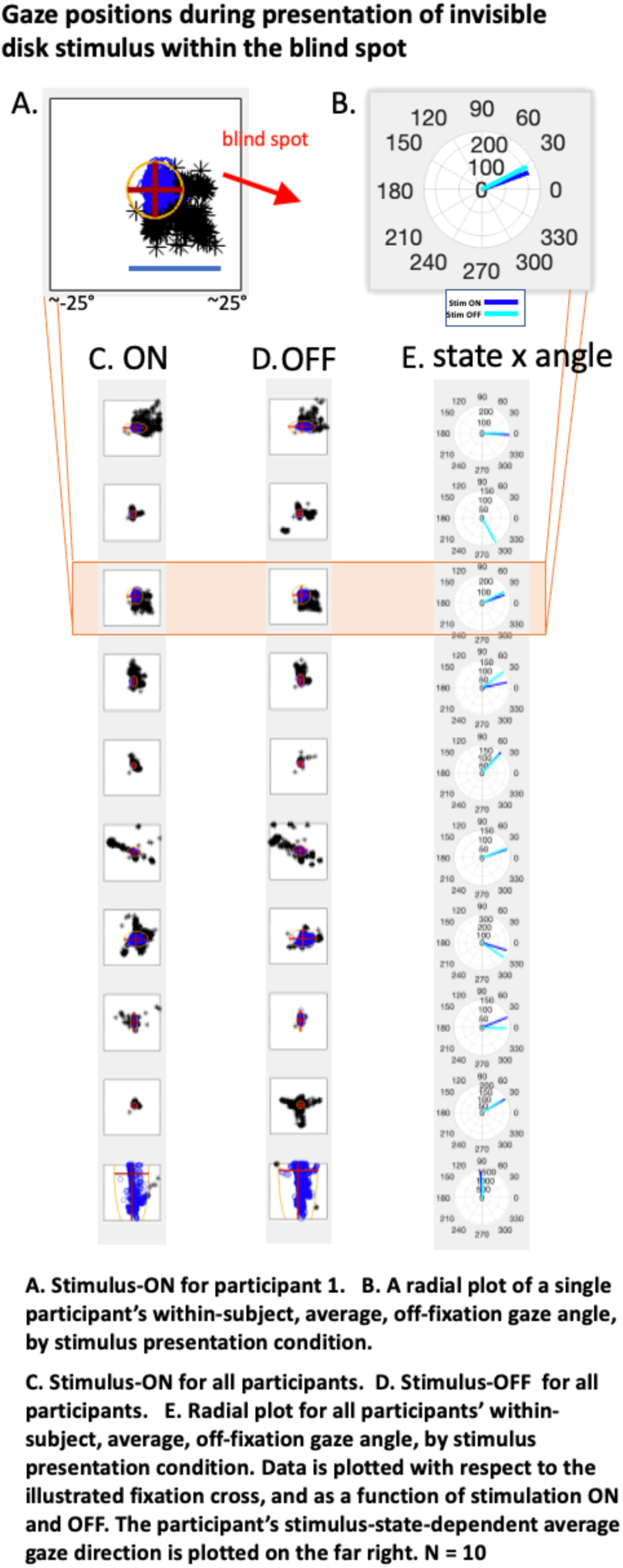
Gaze-position data

We compared eye position data between ON and OFF states, hypothesizing that implicit perception of stimuli within the blind spot might draw a participant’s gaze toward the lower right-side of the visual field, the approximate position of the mapped blind spot for all participants. This follows a body of work on the effects of both implicit and explicit cues on awareness, measuring microsaccades and gaze location (Spering & Carrasco, 2015).

Our results verify that the gaze of fixed-gaze participants is not truly fixed. For example, Fig 19 shows clear spread in gaze position. Further, this participant’s average gaze deflection direction was not to the lower right, toward the blind spot, but to the upper right in both stimulus ON and OFF periods. Looking across participants, during the stimulus ON condition, only two out of the ten participants showed average deviations oriented to the right and below the horizontal meridian, potentially toward the blind spot (Figure 19). Overall, the eye movement data do not show evidence of attentional influence from blind spot stimulation.

## 7. DISCUSSION

The aim of this study was to establish a method to spatially confine stimulation of the retina by employing a laser-based retinal scanning display in psychophysical studies. We developed a hardware setup and demonstrated an efficient method for detailed mapping of the blind spot. We then ran an initial study to investigate implicit processing of blind spot stimulation, to test the validity of our approach, and to find its advantages and disadvantages iteratively.

The disadvantage of this method is that it requires the participant to keep his or her head still for a certain period of time. If the participant looks away from the fixation point, the stimuli disappear. If their posture shifts out of alignment, by even a small degree, the stimuli disappear, or the component colors of the fixation cross split. However, from a different perspective, this disadvantage turned out to be an advantage for enforcing the requirement of participant’s stillness. That is, we were able to turn an important disadvantage of the RSD, its sensitivity to movement, into the useful constraint of gaze-contingency. This stability allowed us to derive participants’ subjective blind spot maps precisely and in a relatively short period of time.

While the EEG and eye movement analyses did not reveal evidence of implicit processes, our measurements of pupil activity showed some positive evidence especially in the longer timescale. Similar to Miyamoto and Murakami (2015) and Gamlin (2007), we found that blue stimuli in the blind spot generate PLR and PIPR (Figures 11 and 15). Measurements of the red control stimuli were marred by a cross-trial artifact. The absence of PLR and PIPR to red light appeared to be obscured by slow modulations left over from previous trials (Figure 16), but an experiment with extended inter-trial recovery periods would be required to confirm this. The two conditions using green ring and field stimuli presented visual stimuli that contained green components that were likely too strong, and should be repeated at lower intensities, in order to detect the effect of ipRGCs inside the blind spot which may be subtle. This may require modification of the laser with respect to green color wavelengths, as the laser’s intensity climbs particularly rapidly through the lowest video levels.

## CONCLUSION

The Retinal Scanning Display allows the collection of detailed functional maps of the visual field and blind spot. It provides an optically cleaner way to study human ipRGCs than with typical displays, by better isolating ipRGC functions from rods and cones. We replicate aspects of previous studies, avoiding scattering artifacts and exploring the parameter space optimized for ipRGC activity. We successfully demonstrated the methodological aspects of applying the device for research. It will be necessary, however, to further test and revise stimulus parameters to make technical improvements aimed at avoiding optical artifacts. This will allow us to further assess the device’s utility and to fully test the hypotheses regarding the biological and functional roles of ipRGCs.

